# Design of peptides with non-canonical amino acids using flow matching

**DOI:** 10.1101/2025.07.31.667780

**Authors:** Jin Sub Lee, Philip M. Kim

## Abstract

The canonical vocabulary of twenty amino acids limits the chemical space available to proteins and peptides. Expanding this vocabulary to hundreds of non-canonical amino acids allows the engineering of proteins with novel function and activity, and is of great interest for the discovery of novel drugs such as macrocyclic peptides. Here we present NCFlow, a flow-based generative model capable of incorporating any arbitrary non-canonical amino acid into a given protein. To supplement sparse training data in the Protein Data Bank, NCFlow is pretrained on millions of small molecule structures and a large set of protein-ligand complexes before finetuning on native non-canonicals found within proteins in the Protein Data Bank. We show that NCFlow outperforms AlphaFold3-based methods in the structure prediction of unseen non-canonical amino acids. We present a peptide design pipeline akin to *in silico* deep mutational scanning, and propose a novel scoring strategy using a combination of deep learning-based and molecular dynamics-based alchemical binding free energy calculations to identify improved peptide variants. We apply the method on four protein-peptide complex test cases, and observe that incorporating non-canonicals can significantly improve binding affinity by up to -7.0 kcal/mol. Thus, NCFlow can be easily integrated into existing protein design platforms to further improve its properties outside of what is capable with standard amino acids.

## 1 Introduction

Proteins typically consist of a polypeptide chain of amino acids, where a standard vocabulary of 20 canonical amino acids build the immense diversity of protein functions necessary for life. However, nature also encodes hundreds of non-proteinogenic non-canonical amino acids (ncAAs) in nature as metabolic intermediates, alterations in the translation pathway, or post-translational modifications of amino acids that allow even greater flexibility in the function and biochemical properties of proteins [1]. Thus, the design of proteins with ncAAs holds significant potential as they can introduce novel functions and capabilities, such as in biocatalysis to develop artificial enzymes, or enzymes with novel reactions not observed in nature [2, 3]. A key area of interest is the design of improved peptide variants with nonstandard backbone or side-chain chemistries. Current approaches to designing such peptides are largely limited to experimental screening via genetic code reprogramming, where modified tRNAs with diversified amino acid payloads are used in conjunction with *in vitro* display technologies such as mRNA display [4]. With the rise in popularity of peptide drugs, and more specifically, macrocyclic peptides containing amino acid modifications, an effective computational approach to nonstandard peptide design would greatly benefit therapeutic drug discovery campaigns.

Protein design with deep learning has been revolutionized in recent years with the development of expressive architectures and rise in data and computing power. AlphaFold2 [5] and its successor AlphaFold3 [6] have facilitated the protein structure prediction boom with near experimental accuracy across many targets and enabled various applications in structure-based protein design [7]. Moreover, the development of diffusion-based generative models [8, 9, 10] and their applications to protein design [11, 12, 13, 14] allows rapid and effective design at unprecedented scale that rivals experimental screening approaches at a fraction of time and cost. However, the adaptation of existing computational protein design tools for nonstandard peptides is non-trivial - AlphaFold2 and RosettaFold-based methods such as RFDiffusion and BindCraft are unable to model ncAAs. Moreover, applying AlphaFold3-based tools such as Boltz1 [15, 16] and Chai1 [17] face another issue since modeling non-standard residues require conditional information (ex. number of atoms per token, RDKit conformer positions) that are specific to each ncAA. Thus, a standard design framework using gradient descent is difficult to apply when the modified residue is not known *a priori*. Concurrent with this work, a method named RareFold and EvoBindRare to design peptide binders with ncAAs was developed [18], which suggests a promising approach using a modified AlphaFold2 architecture and iterative mutational sampling to extend the candidate amino acid pool to 29 ncAAs. Another limitation in modeling ncAAs is the scarcity and bias of ncAA data in the Protein Data Bank, as less than 0.02% of deposited residues correspond to ncAAs. Moreover, a large fraction of these are non-functional, where modified residues such as selenomethionines and selenocysteines are routinely used to aid solving the phase problem in X-ray crystallography. Thus, the fraction of ncAAs that participate in native interactions is severely limited. Finally, scoring ncAA variants is another issue, since most protein-protein and protein-peptide scoring methods [19, 20, 21] only support canonical amino acids, so additional methods to identify high-fitness ncAA variants must be explored.

In this work, we present NCFlow, a model that learns to place any arbitrary ncAA into a given protein backbone. Due to the scarcity of ncAA data, we formulate the task as a single-residue structure prediction problem for which the PDB contains on the order of 10^4^ data points, allowing reasonable training of neural networks. To augment the data, we also employ pretraining on small molecule structures (PubChem [22]) and protein-ligand complexes (Plinder [23]), increasing the available training data by over three orders of magnitude to improve model performance and generalization to ncAAs not found in the PDB. To apply NCFlow in a design setting, we propose an approach similar to deep mutational scanning, where each residue of a given peptide is mutated to a ncAA variant and assessed for improved activity or fitness. We identify improved variants with a combination of uncertainty estimates (pLDDT), deep learning-based affinity prediction (AEV-PLIG [24]), and alchemical relative binding free energy calculations (Alchemical Transfer Method [25]) to identify promising candidates that improve binding affinity to the protein targets. We validate this scoring approach on a comprehensive dataset of experimentally-derived protein-peptide mutational scanning datasets, and observe that across many systems, the proposed combination of deep learning and alchemical methods can effectively isolate improved peptide variants with high precision. We apply the pipeline on four unique test cases including head-to-tail and disulfide-cyclized cyclic peptides, and show that we can obtain peptide variants with ncAAs displaying improved predicted binding affinity across most cases.

## 2 Methods

### 2.1 Data curation

The small molecule dataset is obtained from PubChem3D [22], which uses the OpenEye OMEGA software [26] to generate conformers for >170 million compounds in PubChem. Due to resource constraints, we download a random subset of 13.8 million compounds for pretraining using the single conformer found in PubChem’s FTP site (https://ftp.ncbi.nlm.nih.gov/pubchem/Compound_3D/01_conf_per_cmpd/). We use a 90:10 train:test split and evaluate RMSD on the test set. For protein-ligand finetuning, protein-ligand complexes from Plinderv2 [23] are used, which results in a total of 293,591 complexes after filtering for ligands < 64 heavy atoms and systems for which a *holo* (bound) complex PDB file exists. We set aside 100 randomly selected structures for performance evaluation. The final ncAA-protein dataset is extracted from a snapshot of PDB (dated 2023-07-28) by filtering for the presence of non-canonical residues. This is performed by filtering for all amino acids that are not ‘peptide-linking’ or ‘terminus’ residues as defined in the chemical component dictionary described in PDBeChem (https://www.ebi.ac.uk/pdbe-srv/pdbechem/). We also remove selenomethionine and selenocysteine as these amino acids are routinely used to aid resolving X-ray crystal structures, and therefore are heavily biased in the PDB. We use any ncAA present in the PDB in at least 10 instances to curate the train/validation splits using a random 90:10 split, and test on ncAAs found in less than 10 instances. This results in a total of 38,618, 4,283, and 593 train, validation, and test ncAA examples, respectively, corresponding to 254 unique ncAAs in the train/validation sets, and 231 unique ncAAs in the test set. We note that some of the ncAAs extracted from the PDB exist as ligands in isolation rather than a part of the protein’s polypeptide chain - we chose to use all examples for training due to data scarcity concerns.

### 2.2 Model Training

NCFlow is trained in three stages: first, small molecule conformers in PubChem3D are used to learn general chemical validity given a set of atom types and bond connectivities. Then, the model is finetuned on protein-ligand complexes to learn binding interface biophysics given a protein pocket. Finally, the model is further finetuned on ncAA-protein environments to learn ncAA-specific interactions and placing ncAAs as part of a polypeptide chain. The details for each training stage are described below:

#### 2.2.1 General details

The core framework of NCFlow is based on the I-CFM formulation of flow matching depicted in [27], which we briefly describe here. Flow matching seeks to learn a vector (velocity) field *u*_*t*_ that defines an ODE, where the solutions to this ODE are defined by a flow *ψ*_*t*_. X is defined as the trajectory that follow the vector field from *X*_0_ to *X*_1_, where *X*_0_ *∼ p* and *X*_1_ ∼ *q*, where *p* and *q* are the prior and data distributions, respectively. Thus, the ODE is defined by:

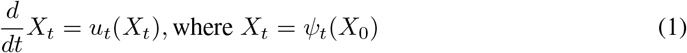

For generative modeling, we want to build a flow *ψ*_*t*_ that yields some probability path *X*_*t*_ ∼ *p*_*t*_ that transforms a sample from a simple source distribution (typically *X*_0_ ∼ *𝒩* (0, 1)) to one from the complex target distribution *X*_1_ ∼ *p*_*data*_(*X*), and solve the ODE determined by the vector field *u*_*t*_ from *t* = 0 to *t* = 1. There are infinitely many choices to define *ψ*_*t*_ and thus *p*_*t*_, but the simplest formulation is to construct a set of linear conditional paths *p*_*t*_(*X* |*X*_1_) for every training data *X*_1_ ∼ *q*, resulting in the following marginal:

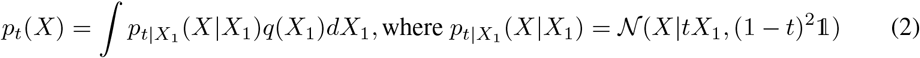

which results in a simple expression for *X*_*t*_ that takes a linear combination of *X*_1_ and *X*_0_ dependent on *t*:

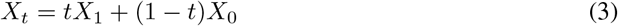

and a regression target of *u*_*t*_ as follows:

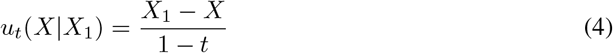

Rather than conditioning on just *X*_1_, [27] reports a more flexible approach to define the probability path for arbitrary source distributions by independent coupling of source and target points (referred to as I-CFM), resulting in the following expressions:

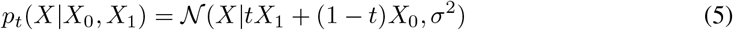

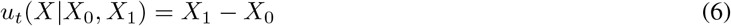

Equipped with this, we can define an objective function to train a neural network parameterized by *θ* to approximate *u*_*t*_:

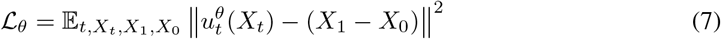

where *t* ∼ *U* [0, 1], *X*_0_ ∼ *p, X*_1_ ∼ *q*.

For sampling, we initalize the trajectory with *X*_0_ ∼ *p* and apply a numerical ODE solver such as Euler’s method defined below:

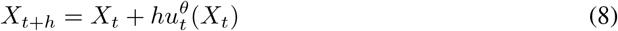

where *h* is a stepsize hyperparameter indicating fixed timesteps *t* ∈ [0, 1]. We use 10 timesteps for all cases, and found no performance improvements with increased timesteps. Since we do not use an equivariant architecture (see *Model Details*), we use data augmentation at every iteration by randomly rotating and translating all atoms with a standard deviation of 1A across all stages of training. We also use exponential moving average of model parameters with a decay of 0.999 per iteration, and gradient clipping with norm 1.0. All models are trained on 4 NVIDIA A100s (40GB) with DistributedDataParallel.

#### 2.2.2 Stage 1: small molecule pretraining

The initial pretraining stage on PubChem3D is rather straightforward, as the model simply learns to generate the full conformer given the atom types and bond connectivity graph. The conformers are centered at mean zero, and the model is trained with effective batch size of 32 and learning rate at 2*e* − 4. The model was trained for approx. 7 days, corresponding to 10 epochs.

#### 2.2.3 Stage 2: protein-ligand finetuning

For protein-ligand data, the model takes in conditional pocket information defined by the 128 atoms closest to the ligand’s centroid. Only the ligand is noised, and the loss is propagated from just the ligand atoms to train the model. The pocket and ligand coordinates are centered on the ligand’s centroid to provide coarse information on the ligand coordinates - we note that this is not ideal if the desired task is protein-ligand docking or small molecule design. However, NCFlow is trained ultimately for single-residue ncAA structure prediction for which some conditional information (ex. backbone atom coordinates) exist, and therefore allowed such biases to be present in this training stage. The model was trained with effective batch size of 16 and learning rate 2*e* − 5. The model was trained for approx. 7 days, corresponding to 15 epochs.

#### 2.2.4 Stage 3: ncAA-protein finetuning

To train on ncAA-protein environments, the pocket definition was expanded to 200 atoms to allow greater context when predicting the ncAA. The prior distribution is set as a Gaussian distribution with mean on the residue’s CA coordinate. We also introduce local side-chain flexibility of the pocket residues by introducing Gaussian noise centered on the ground truth coordinates with standard deviation 0.2A, which then the model learns to locally repack neighboring side-chains while predicting the ncAA structure. Following [28], residues for local side-chain repacking were selected if any atom of the residue lies within 3.5A of any ncAA side-chain atom, which may increase the number of pocket atoms - we cropped this to a maximum of 256 atoms for training efficiency. Backbone atoms (including the carbonyl oxygen) remain fixed throughout training. At every training iteration, we sample a random structure corresponding to a ncAA to limit biases of certain ncAAs in the PDB. The model is trained with effective batch size of 4 and learning rate 2*e* − 6 for approx. 7 days.

Since the model does not explicitly enforce equivariance, we observe that chirality issues arise in some samples. We resolve this post hoc by discarding generated samples that do not exactly match the configuration of the chiral centers denoted in the input ncAA SMILES string (using RDKit’s FINDMOLCHIRALCENTERS function). A more direct approach to resolve chirality issues such as Feynman-Kac steering [29] used in Boltz1 [15] may be beneficial, but the low memory and runtime requirements of NCFlow (<4GB VRAM and 5 seconds per sample for on a NVIDIA RTX3060) does not necessitate this. Moreover, we observe that 8 samples per ncAA is sufficient for obtaining ncAAs with correct chirality in almost all cases.

### 2.3 Model architecture and featurization

The model is based on AlphaFold3’s Pairformer and Atom Transformer modules implemented in Boltz1. We remove the tokenization scheme used in AlphaFold3, and rather use an atom-level representation throughout the model. The node features consist of the atom types, timestep, noised coordinates, and a binary mask indicating the target atoms to predict (ex. ligand atoms for Pub-Chem/Plinder, ncAA for PDB). For the final finetuning stage on ncAA-protein environments, we also include a mask to indicate the sidechains for repacking. The pair features are the pairwise Euclidean distances and one-hot-encoded bond connectivities, which contains five labels indicating single, double, triple, aromatic, or no bonds between two atoms. The model contains 16 Pairformer layers and 4 Atom Transformer layers with single and pair hidden dimensionality of 128 and 64, respectively, resulting in 8.7 million trainable parameters. The confidence model is a down-sized version of the main model with 6 Pairformer layers and 1 Atom Transformer layer. The confidence model is trained to predict per-atom LDDT scores using a similar ‘diffusion rollout’ scheme adopted by AlphaFold3, where the main model is frozen and candidates are sampled using 10 timesteps which are then scored using the confidence model.

### 2.4 Design pipeline

To design peptides with ncAAs, we use *in silico* deep mutational scanning in conjunction with binding affinity estimators to identify variants that increase binding affinity to a given target. The details are described below in their respective sections.

#### 2.4.1 Mutational scanning

Deep mutational scanning is a standard experimental technique for protein engineering, where proteins are mutated at every position with all 20 canonical amino acids to produce a library of variants, which then undergo a selection process to filter high fitness variants and sequencing for identification. We follow a similar approach here, where all positions of the peptide are mutated with a pool of ncAAs, which are assessed through *in silico* binding affinity estimators to identify variants that increase binding affinity. The pool of ncAAs are obtained by curating synthesizable ncAAs from WuXi AppTec, a peptide synthesis CDMO company. Specifically, we download all ncAAs linked to Fmoc, have a related parent canonical amino acid, and are parseable with RDKit, resulting in a total of 939 ncAAs. To reduce combinatorial complexity, we only mutate each residue to a related ncAA according to its parent amino acid, which can range from 3-379 variants (except isoleucine, which does not have any variants after these filters). The exact number of ncAAs per parent amino acid is listed in Table S1. Using the curated pool of ncAAs, we generate a large pool of structural variants at each position for each protein-peptide complex test case.

#### 2.4.2 Binding affinity estimation

The presence of ncAAs makes it difficult to use any existing protein-peptide binding affinity prediction tools as they typically can only encode canonical residue identities. Thus, we explore using protein-ligand binding affinity prediction methods which are more flexible in the way that the ligands are encoded (ex. atom types and bond graph). Specifically, we use a deep learning-based tool called AEV-PLIG [24], which is trained on PDBBind2020 [30] and other augmented datasets to predict protein-ligand binding affinities closer to FEP+ calculations. We also note that PDBBind contains peptide ligands, so we reasoned that the model can generalize to predict ncAA-containing protein-peptide binding affinities. To protonate the peptides we use the hydride package within Biotite [31], which can then be processed by AEV-PLIG to predict *pK*_*d*_ values.

We also use an alchemical relative binding free energy prediction tool called Alchemical Transfer Method (ATM) [25], which runs MD ensembles with an alchemical coordinate transformation that translates one ligand from the binding pocket to the solvent, and another ligand from the solvent to the binding pocket. By calculating the free energy differences between the alchemical states, we can measure relative binding affinity of two ligands of interest. To prepare the system for ATM, we solvate the system with 10nm padding at pH 7.0, and use Antechamber to prepare the ligands with the GAFF2 force field and AM1-BCC charge model [32]. The wild-type peptide is left inside the binding pocket, and the peptide ncAA variant is displaced from the binding site by a coordinate translation of magnitude 25A along the vector from the target protein’s centroid and the ligand binding site. We use the first three CA atoms for aligning the peptide reference atoms and positionally restrain all CA atoms of the protein target. The binding site restraints are set to all CA atoms for both the peptide and receptor. The system is simulated with 2fs timesteps using 8000 steps per sample for 200 samples across 22 replicas (alchemical states), corresponding to a 71ns MD ensemble per variant, or an average of 12 hours of runtime on a single NVIDIA A100 (40GB). We use the Acellera HTMD package [33] for system preparation and the Quantumbind-RBFE [34] pipeline to run ATM, using the Amber ff14sb force field [35] to parameterize the target protein. We experimented with using the AceFF 1.0 neural network potential to run NNP/MM simulations as in QuantumBind-RBFE, but the computational cost of using the NNP on longer ligands such as peptides made it infeasible to run in high-throughput. We tested using NNP/MM with ATM on three variants on a protein-peptide complex and observed comparable values with the GAFF2 force field at 10X computational cost - while inconclusive, this requires further investigation that we leave for future work. We also observe that some simulation runs fail due to exploding energy or coordinate issues, which can be a consequence of improperly defined constraints, initial positions, or simulation parameters (timestep, etc.). We observed that this issue is exacerbated with increased timesteps at 4fs with hydrogen mass repartitioning to 4 amu. Since molecular dynamics simulations often require system-specific optimization for accurate modeling, it may be possible that the current set of hyperparameters causes some simulations to explode. Thus, an optimized ATM protocol applicable to all protein-peptide system is a direction of future work.

## 3 Results

First, we evaluated the respective pretrained models on PubChem3D and Plinder on their ability to recover the ground truth small molecule conformations (Figure 2A). Since PubChem3D consists of small molecules in isolation, we use Kabsch-aligned ground truth and sample structures to calculate RMSD, and observe that the model generates small molecule conformers with high fidelity (Supplementary Figure 1A). For Plinder evaluation, we observed that using standard RMSD sometimes inflates the values when the ligand contains symmetric groups, and the model predicts two symmetric groups in an alternative yet chemically equivalent form than the ground truth complex. Thus, we use a symmetry-corrected RMSD [36] to evaluate performance, which computes all isomorphic graphs of the ligand (which simply reorders the small molecule atoms) and identifies one corresponding to the lowest RMSD. We observe that the protein-ligand finetuned model performs reasonably well with a mean RMSD of 2.23Å, with 62% and 33% of conformers displaying RMSD < 2.0Å and < 1.0Å, respectively. Moreover, the generated conformers are realistic and exhibit various reasonable interactions with its target protein (Supplementary Figure 1B). However, we emphasize that this metric is not comparable with blind protein-ligand docking because we provide coarse information on the ligand’s binding pocket, and the test set is randomly selected, which may result in leakage issues. However, we did not pursue more robust evaluation as protein-ligand docking is not the task of interest in this work.

**Figure 1:**
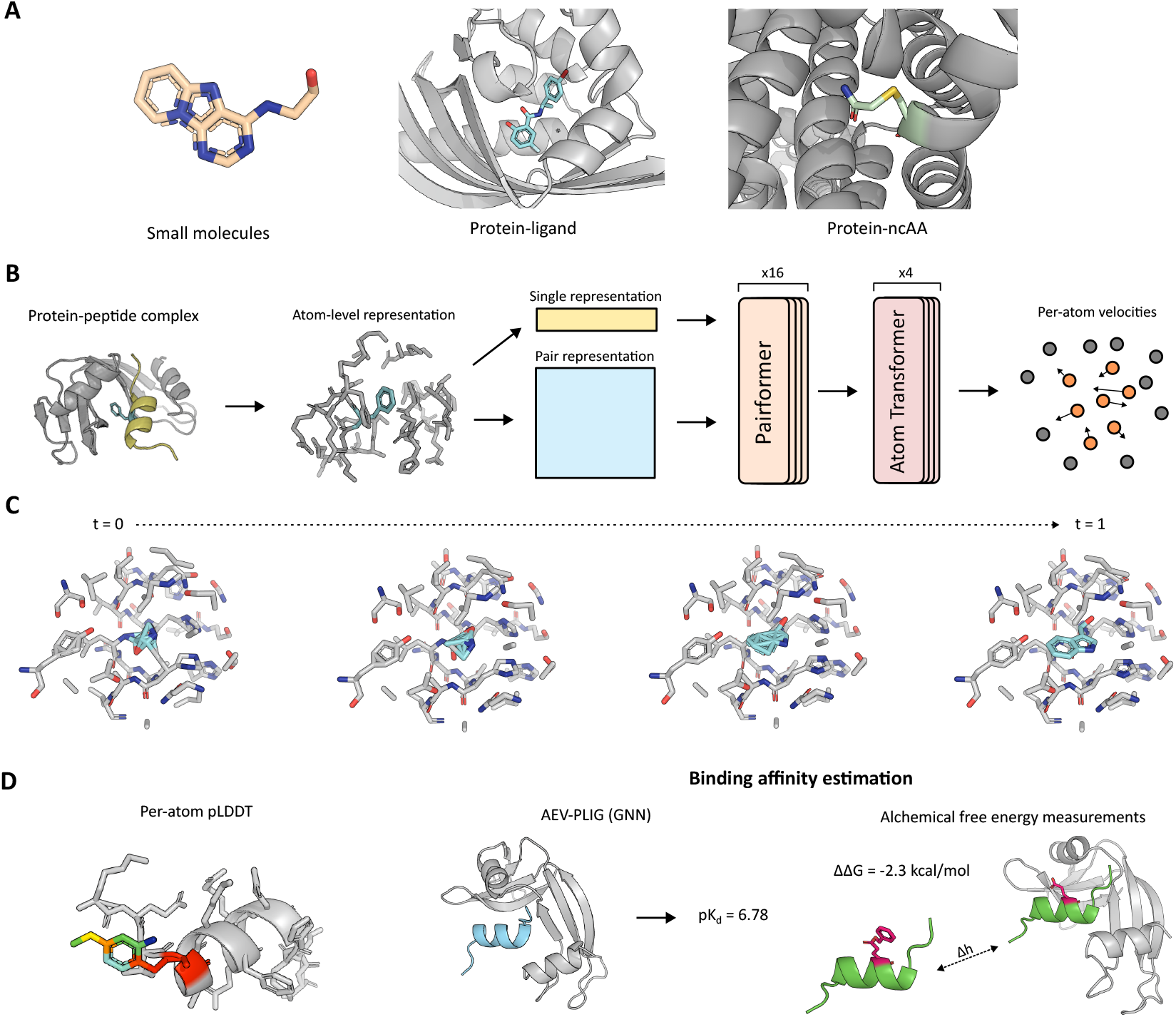
Overview of NCFlow. A) NCFlow is trained from three datasets: 1. PubChem3D, containing low-energy conformers of small molecules, 2. Plinder, a curated dataset of protein-small molecule complexes, and 3. a dataset of ncAA-protein environments found in the Protein Data Bank. B) NCFlow simply takes in a protein-peptide complex and the chemical description of a ncAA (atom types and bond connectivity), and predicts the 3D conformation of the ncAA in the given residue and pocket. C) An example sampling trajectory of a ncAA using NCFlow from *t* = 0 to *t* = 1. D) To enable design, we rely on three filters: per-atom pLDDT predicted by a confidence module, a deep learning-based protein-ligand binding affinity predictor named AEV-PLIG [24], and MD-based relative binding free energy calculations via an alchemical coordinate transformation Δ*h* [25].

**Figure 2:**
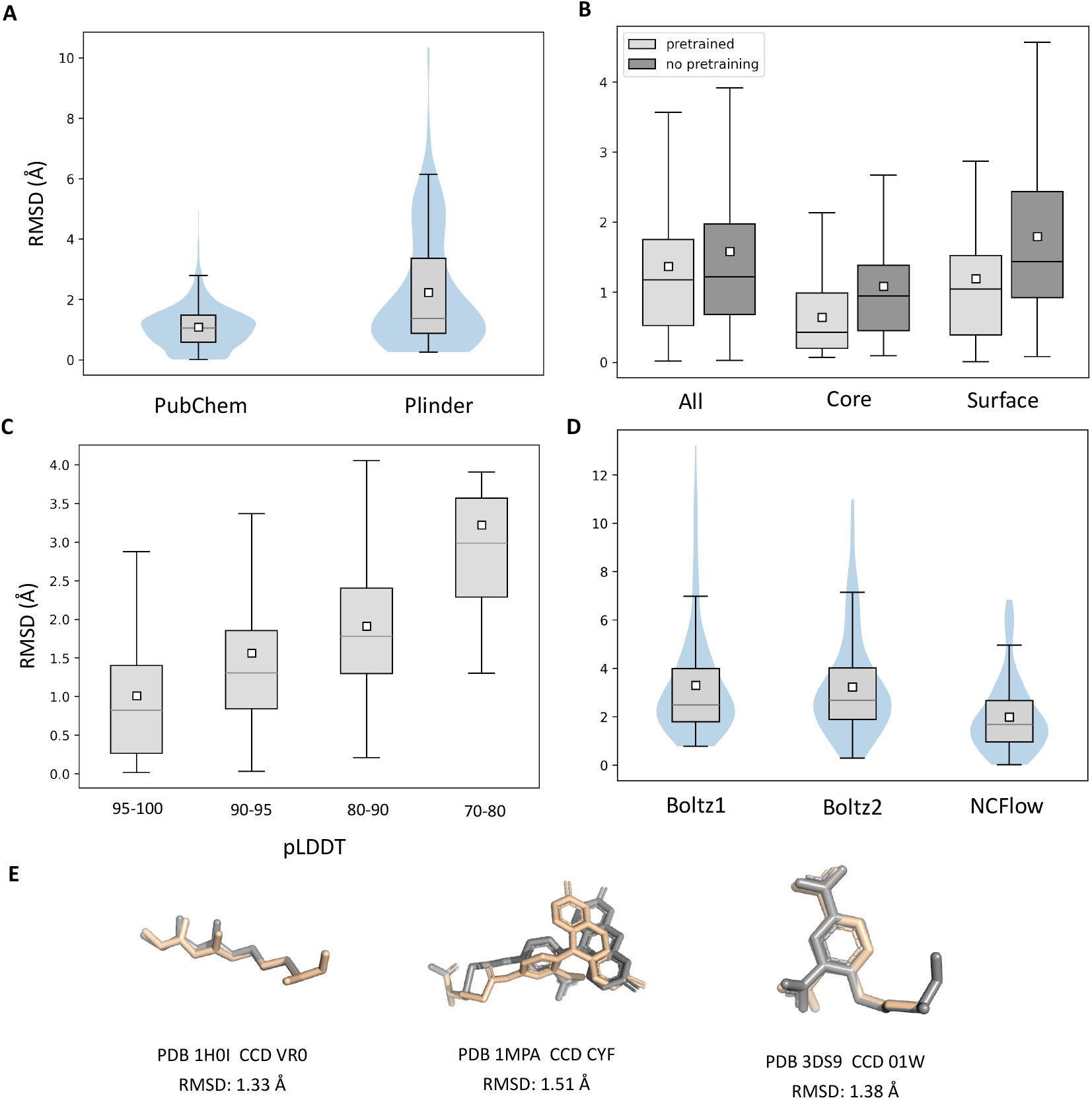
Performance evaluation of NCFlow. A) RMSD plots of PubChem and Plinder pre-trained models. B) RMSD evaluation of NCFlow trained from scratch and finetuned from the PubChem/Plinder pretrained model. Across all, core (> 35 CA atoms within 12Å), and surface (< 20 CA atoms within 12Å) residues, the finetuned model outperforms the model trained from scratch.C) Binned pLDDT and RMSD distributions show that high-confidence samples can be used to filter out poor predicted conformers. D) NCFlow outperforms Boltz1 and Boltz2 in single-residue ncAA prediction with an mean RMSD difference of 1.86Å and 1.81Å, respectively. E) Randomly selected ncAA predictions by NCFlow, with ground truth in gray and generated conformers in beige. Note that the white square box in all boxplots represent the mean value of the distribution.

We then tested whether the PubChem3D and Plinder pretraining tasks improves performance on prediction of single-residue ncAA structures given its protein pockets (Figure 2B). We observe that no pretraining attains strong performance with a mean RMSD of 1.58Å across all test set ncAAs, with lower RMSDs observed for core residues (1.08Å) than surface ones (1.79Å) due to increased contextual information and therefore rigidity of the ncAA in the protein pocket. Pretraining on PubChem and Plinder further decreases RMSD across the test set by 0.21, 0.14, and 0.18Å in all, core, and surface residues, respectively, suggesting the effectiveness of pretraining on small molecules for ncAA structure prediction. We then analyzed the intrinsic uncertainty estimates (pLDDT) predicted by the confidence model for ranking predictions (Figure 2C), and observed a strong negative relationship between pLDDT and RMSD. This suggests that high-confidence samples ranked by pLDDT can be used to filter out poor samples generated by NCFlow.

We also report comparisons to Boltz1/2 [15], an open-source implementation of AlphaFold3 that is capable of predicting protein structures with ncAAs (Figure 2D). Direct performance comparison is not possible since AlphaFold3-like models are trained for the much more difficult task of general biomolecular structure prediction and the non-ncAA protein residues are co-folded with the ncAAs, while NCFlow considers only the pocket and neighboring side-chains and assumes the protein backbone to be fixed. Thus, poor predictions of the overall protein scaffold by Boltz1/2 may lead to large deviations in RMSD. We ameliorate this issue by 1. superimposing the predicted ncAA backbone atoms (N, CA, and C) by Boltz1/2 with the ground-truth ncAA backbone atoms prior to RMSD calculation, and 2. running NCFlow in single-chain mode by isolating the chain containing the ncAA. We observe that Boltz1 and Boltz2 performs significantly worse than NCFlow for single ncAA structure prediction with mean RMSDs of 3.29Å, 3.24Å, and 1.43Å respectively, suggesting that NCFlow predicts more accurate ncAA conformations over AlphaFold3-based methods. We report a few ground-truth and predicted ncAA structures from the test set in (Figure 2E), and observe that the predicted ncAA structures are structurally valid and align closely with the ground-truth conformations.

To apply NCFlow for engineering peptides with ncAAs, we use *in silico* deep mutational scanning with virtual screening via computational binding affinity predictions to identify peptide variants with improved predicted binding affinity (Figure 3A). We start with a bound protein-peptide complex, and isolate the pocket atoms based on the CA atom of the peptide residue of interest. Then, we extract a pool of ncAAs based on the wild-type parent amino acid in the form of SMILES strings, which can be used to extract atom types and bond connectivity information. NCFlow takes in the pocket structural data and ncAA atom type and bond information to replace the parent canonical amino acid to the ncAA variant, resulting in a protein-peptide complex containing the selected ncAA. We predict 8 samples per variant, and select the peptide conformer with correct chirality and highest pLDDT for downstream analysis. This process is iterated over all residue positions and the entire ncAA pool to obtain hundreds of protein-peptide complex variants, each containing a unique peptide-ncAA variant bound to the target protein. These variants are filtered to exclude low-confidence predictions by a pLDDT filter of > 90, and AEV-PLIG [24] is used to predict binding affinity of the wild type peptide and the ncAA-containing variant, which has been shown to exhibit state-of-the-art performance and outperforms ipTM confidence scores of Boltz-1x across most metrics [37]. We exclude all variants with a AEV-PLIG predicted Δ*pK*_*d*_ < 0 based on the wild-type predicted *pK*_*d*_, and select the top 50 variants by highest Δ*pK*_*d*_ for further evaluation.

**Figure 3:**
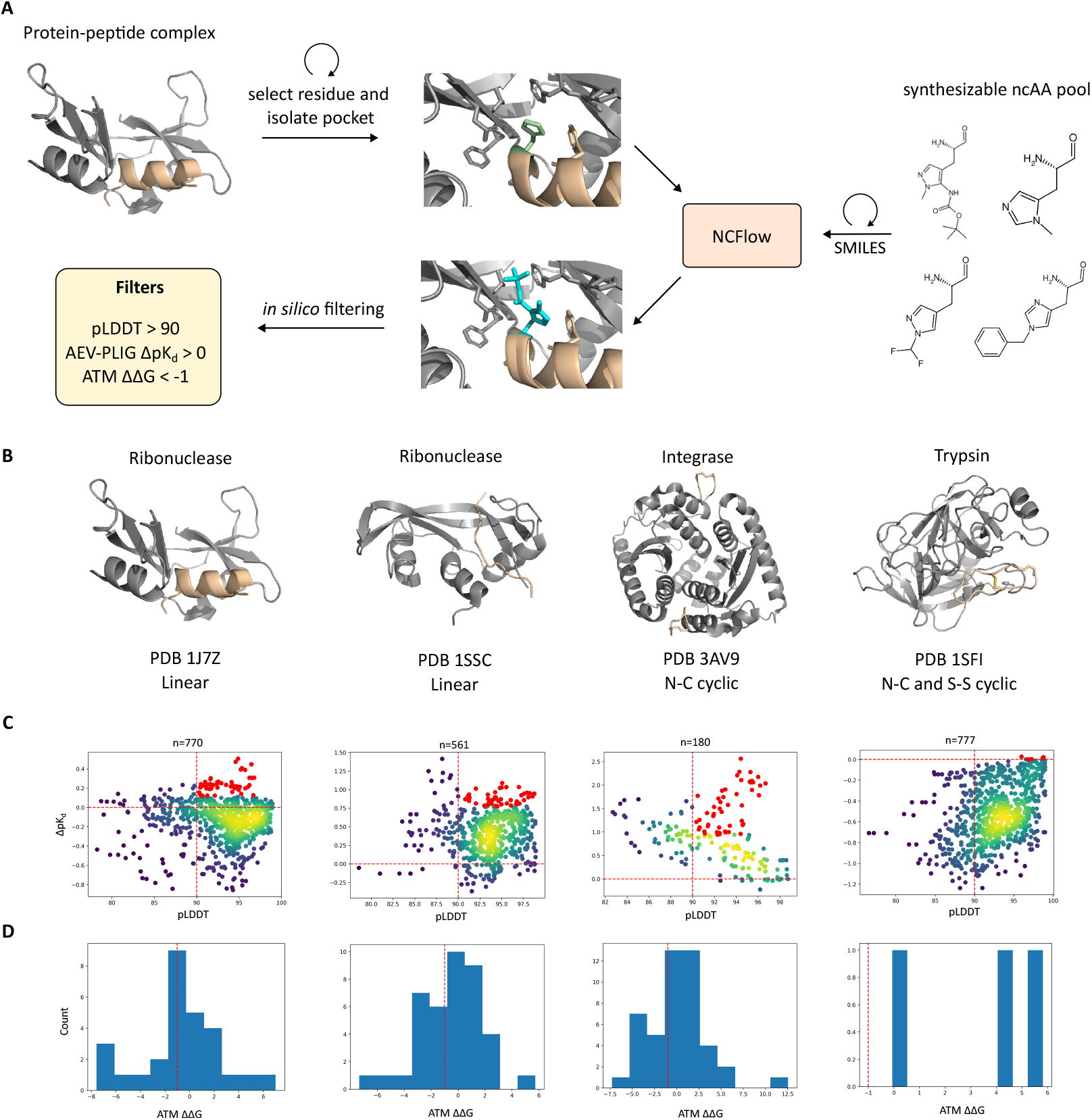
Design of peptide-ncAA variants using NCFlow. A) First we take a bound protein-peptide complex, select a peptide residue, and isolate its pocket environment. NCFlow receives the pocket information and ncAA data in the form of SMILES from which the atom types and bond connectivity is extracted. NCFlow predicts the 3D conformation of the ncAA into the given residue, which is scored using a set of filters for prediction accuracy and binding affinity. This is repeated across all peptide residues and ncAAs. B) Selected protein-peptide test cases, ranging from helical linear peptides (1J7Z) to head-to-tail and disulfide-cyclized peptides (1SFI). C) pLDDT and AEV-PLIG Δ*pK*_*d*_ distributions of all ncAA variants in each test case. Yellow and blue indicate high and low density regions, respectively, and red indicate the top 50 selected variants by Δ*pK*_*d*_. Vertical and horizontal red dashed lines indicate pLDDT = 90 and Δ*pK*_*d*_ = 0, respectively. D) ATM ΔΔ*G* measurements of top 50 variants for each test case.

Since peptide-ncAA variants are costly to synthesize, we further validate these variants by measuring the relative binding free energy using the Alchemical Transfer Method [25], an alchemical free energy method that estimates binding affinity by alchemical coordinate transformations between two ligands (see *Methods*). Thus, we obtain tens of variants per protein-peptide complex that are of high-confidence by pLDDT, and are predicted to increase binding affinity by two orthogonal methods - a deep learning-based model (AEV-PLIG) and a more rigorous MD-based alchemical relative binding free energy measurement (ATM). We first tested AEV-PLIG on long ‘peptides’ in the PDBBind-PP dataset not used for training, and observed a moderate Spearman correlation of 0.39 on absolute binding affinity measurements. This suggests that AEV-PLIG can indeed be used for assessing protein-peptide complexes despite being trained on smaller chemical ligands and short peptides. On a test run on 28 peptide variants containing a relatively smaller pool of ncAAs from CycPeptMPDB [38] on the target PDB 1SSC, we observe a negative Spearman correlation of -0.62 between AEV-PLIG predicted Δ*pK*_*d*_ and ATM ΔΔ*G* (Supplementary Figure 2), suggesting that the two deep learning-based and alchemical methods are moderately aligned on variants that constitute strong and poor binders.

To evaluate the proposed scoring method on known protein-peptide complexes, we first curated multiple experimental binding affinity datasets from literature. We obtained four sets of protein-peptide complexes containing multiple mutations to canonical amino acids and corresponding binding affinities [39, 40, 41], which allows us to assess the effectiveness of the AEV-PLIG/ATM scoring method on experimentally validated affinities. We generated 3D structure for each variant with either PyMOL’s mutagenesis tool (which simply finds the lowest-energy rotamer), or AlphaFold2 using high-confidence (pLDDT > 90) and low-RMSD (<1.5A) predicted structures. We predicted the binding affinity of each variant using AEV-PLIG, and relative ΔΔG measurements using the ATM protocol compared to a given reference peptide (by selecting the wild-type or main peptide scaffold used by each paper), and report the results in Supplementary Figure 5 and Supplementary Table 5. Interestingly, we observe that correlations with binding affinity were often weak or insignificant in most cases, with the exception of ATM on Mdm2-p53 which exhibits a remarkable 0.76 correlation with its experimental binding affinity. Moreover, AEV-PLIG exhibits a moderate correlation of -0.43 and -0.48 with Htra1-PDZ and cetuximab-meditope complexes, though with p > 0.05 due to insufficient number of samples. However, if we apply the cutoff of AEV-PLIG Δ*pK*_*d*_ < 0 and ATM ΔΔ*G* < -1.0 kcal/mol and analyze the confusion matrices, both methods show strong performance in distinguish stronger vs weaker binders compared to the reference peptide, with ATM often exhibiting higher accuracy and precision than AEV-PLIG across the four test complexes. Remarkably, combining both methods with their respective cutoffs results in a precision of 1.00 across all samples, suggesting that applying the two orthogonal methods for binding affinity prediction is a promising approach for identifying peptide variants with increased binding affinity. We then tested the protocol on a deep mutational scanning dataset of two protein-peptide complexes with 41 canonical and non-canonical amino acids reported in [42], and report the results in Supplementary Figure 6. The PUMA peptide showed a promising yet weak negative correlation of -0.19 with precision of 0.36 and recall 0.79 using the Δ*pK*_*d*_ < 0 cutoff (given a 29-71 positive-negative imbalanced dataset), but running ATM on PUMA almost always failed due to the length of the peptide (35aa) causing simulation errors. On the other hand, the CP2 peptide failed to exhibit any notable correlation with neither AEV-PLIG or ATM on 150 selected variants, suggesting that the scoring method works on a system-specific basis and reiterates the need for more robust scoring methods beyond existing deep learning-based and alchemical approaches. We also note that the aforementioned dataset calculates ΔΔ*G* using relative enrichment of DNA sequencing reads calibrated to known binding affinities and therefore may not capture true ΔΔ*G* upon binding.

We ran the design pipeline on a total of four test cases, ranging from standard linear helical peptides to head-to-tail or disulfide-bridged cyclic peptides (Figure 3B). We plot the pLDDT with AEV-PLIG Δ*pK*_*d*_ for all variants to analyze the variant distribution for each test case (Figure 3C). Filtering for high-confidence samples removes 22.4% of the samples across all test cases, suggesting that most generated samples are predicted with low RMSD. Moreover, 47.1% of the variants exhibit a Δ*pK*_*d*_ > 0, indicating that more than half of the ncAA variants are predicted to decrease binding affinity. We measured the Spearman correlation between pLDDT and Δ*pK*_*d*_ and observe large variance ranging from -0.40 (3AV9) to 0.39 (1SFI), suggesting that internal confidence metrics cannot be used for binding affinity estimation. For some test cases (PDB 1J7Z and 1SFI), we observe that most of the variants have a predicted Δ*pK*_*d*_ < 0 with 25.8% and 0.64%, respectively, suggesting that most ncAA variants do not improve binding affinity relative to the wild-type peptide. Interestingly, the head-to-tail and disulfide-cyclized peptide in PDB 1SFI contain only 5 variants that minimally improve binding affinity. This can be explained by the tight binding affinity of the native cyclic peptide to trypsin - its cognate receptor - with an experimentally measured binding affinity of 0.1nM (and in AEV-PLIG’s PDBBind training set), while the other test cases are measured and/or predicted to exhibit micromolar affinity. Thus, it may be increasingly difficult to identify variants that further improve binding affinity of strong binders, especially with just single mutants.

Finally, we ran the top 50 variants ranked by AEV-PLIG Δ*pK*_*d*_ for each test case with ATM to further computationally validate these peptide-ncAA variants (Figure 3D). We observed that 35.0% of samples exhibit ATM ΔΔ*G* < -1.0 kcal/mol, indicative of variants that are predicted to increase binding affinity by both binding affinity prediction tools. Interestingly, the 3 ncAA variants of PDB 1SFI where ATM successfully ran yielded no variants with ATM ΔΔ*G* < -1.0 kcal/mol, so we removed this complex from further downstream analysis. We measured the correlation between AEV-PLIG Δ*pK*_*d*_ and ATM ΔΔ*G* of the top 50 variants across all test cases, and observed poor correlation with no statistical significance (Supplementary Figure 3), suggesting that the two methods do not necessarily agree on the ranking of high-affinity binders perhaps due to limited dynamic range.

We visualize some of the top variants per test case in Figure 4. We observe that the variants predicted to increase binding affinity exhibit either increased noncovalent interactions with pocket residues or increased polar interactions with the solvent. For instance, the ALA6 variant of PDB 1J7Z (Figure 4A, middle) contains a 2-hydroxy-4-methoxybenzaldehyde group linked to the peptide nitrogen atom that induces polar interactions with the solvent, suggesting it may stabilize native binding interactions with the target protein. Moreover, the PHE120 variants of PDB 1SSC (Figure 4B) exhibit increased side-chain interactions with neighboring pocket residues in distinct ways: the left variant is linked to an iodine atom that exhibit polar interactions with the neighboring H12 residue, while the right variant contains a chlorine atom in the same position for polar interactions in addition to a C-N group that extends into a pocket consisting of polar oxygen atoms of T45, D83, and S123 residues of the target protein for increased hydrogen-bonding interactions. The ALA2 variant of PDB 3AV9 contains two solvent-exposed polar fluorine atoms for increased solvent interactions, and the ASP5 variant appears to exhibit *π* − *π* interactions with the protein’s Y99 residue (Figure 4C). Thus, we show that the design workflow can generate ncAA variants for protein-peptide test cases that are biophysically plausible to increase binding affinity to its target protein.

**Figure 4:**
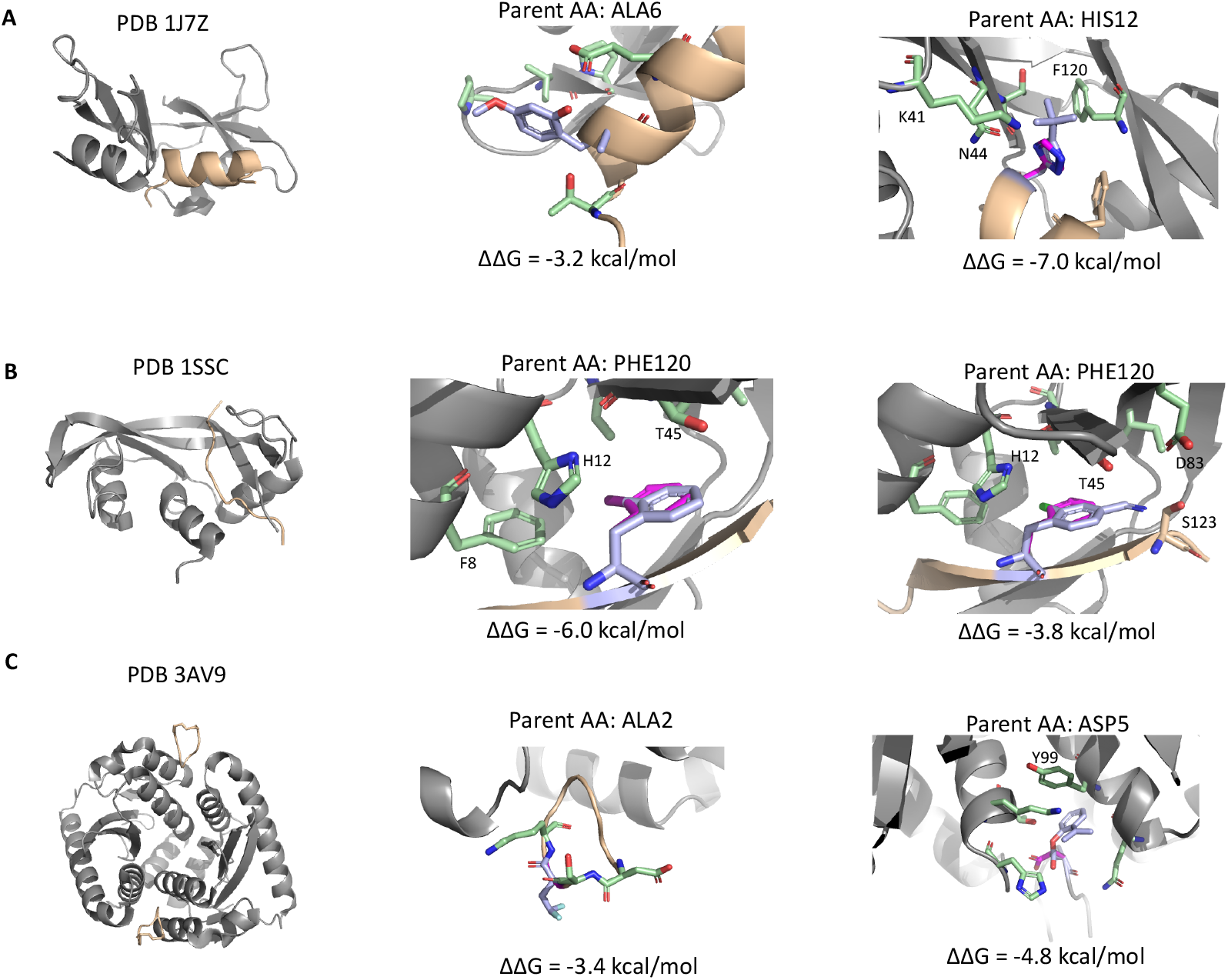
Selected ncAA variants with improved binding affinity for each test case. The ground truth pocket is colored in gray with selected interacting pocket residues in green, the wild type residue to be replaced in magenta, and the ncAA variant in blue.

## 4 Conclusion

In this work, we present a novel framework to incorporate ncAAs into protein-peptide complexes to improve binding affinity. We developed NCFlow, a flow matching generative model that learns to predict the 3D conformations of any input ncAA within a given protein pocket, and apply the model to generate hundreds of peptide-ncAA variants that are scored with deep learning and alchemical methods to predict binding affinity. We evaluate the proposed scoring methods, originally tailored for protein-ligand complexes, on known protein-peptide binding affinities and observe that in some systems, we attain strong performance with precision 1.0, but does not universally apply across all protein-peptide systems. We show that the framework can be used to generate novel peptide-ncAA single mutants that are predicted to increase binding affinity, presenting a novel method of protein engineering that can supplement existing *de novo* protein design workflows.

We envision many possible avenues of future work, the most notable of which is experimental characterization of these peptide-ncAA variants. Though computational binding affinity prediction has progressed significantly with the development of deep learning models and application of improved force fields such as neural network potentials (NNPs) [34], they are imperfect proxies that are often not universally applicable to all systems and experimental validation is still required to validate the effectiveness of NCFlow’s design approach. As such, a more thorough evaluation of using computational approaches such as ATM for protein-peptide binding free energy calculation is required, but the intensive computational resources required by alchemical methods such as ATM limits extension to large sets of experimentally characterized protein-peptide affinities. As more efficient and accurate computational methods for binding affinity prediction are developed - perhaps through extension of Boltz2 to protein-peptide complexes - we believe a design pipeline as the one depicted in this work will become increasingly more useful. Another direction of future work is the identification of a smarter pool of ncAAs that are functionally relevant, as we observed that many synthesizable variants available on Wuxi AppTec carried side-chain protecting groups that are less likely to participate as favorable interactions. Additionally, we only focus on single mutants to reduce computational complexity, yet the workflow can easily be extended to incorporate additional ncAA variants that may further improve binder properties, not limited to just binding affinity. Ultimately, a peptide design pipeline that can simultaneously incorporate multiple ncAAs and co-folds the protein-peptide complex in the style of AlphaFold3 would be greatly beneficial, which may be possible as more ncAA-containing proteins are deposited in the Protein Data Bank and more expressive and performant models are developed.

## 5 Acknowledgements

We thank the many open-source codebases that this work was based on, including PyTorch, Boltz, QuantumBind-RBFE and AToM-OpenMM. We also would like to thank Digital Research Alliance of Canada for computing resources.

## Supplementary Information

**Figure S1:**
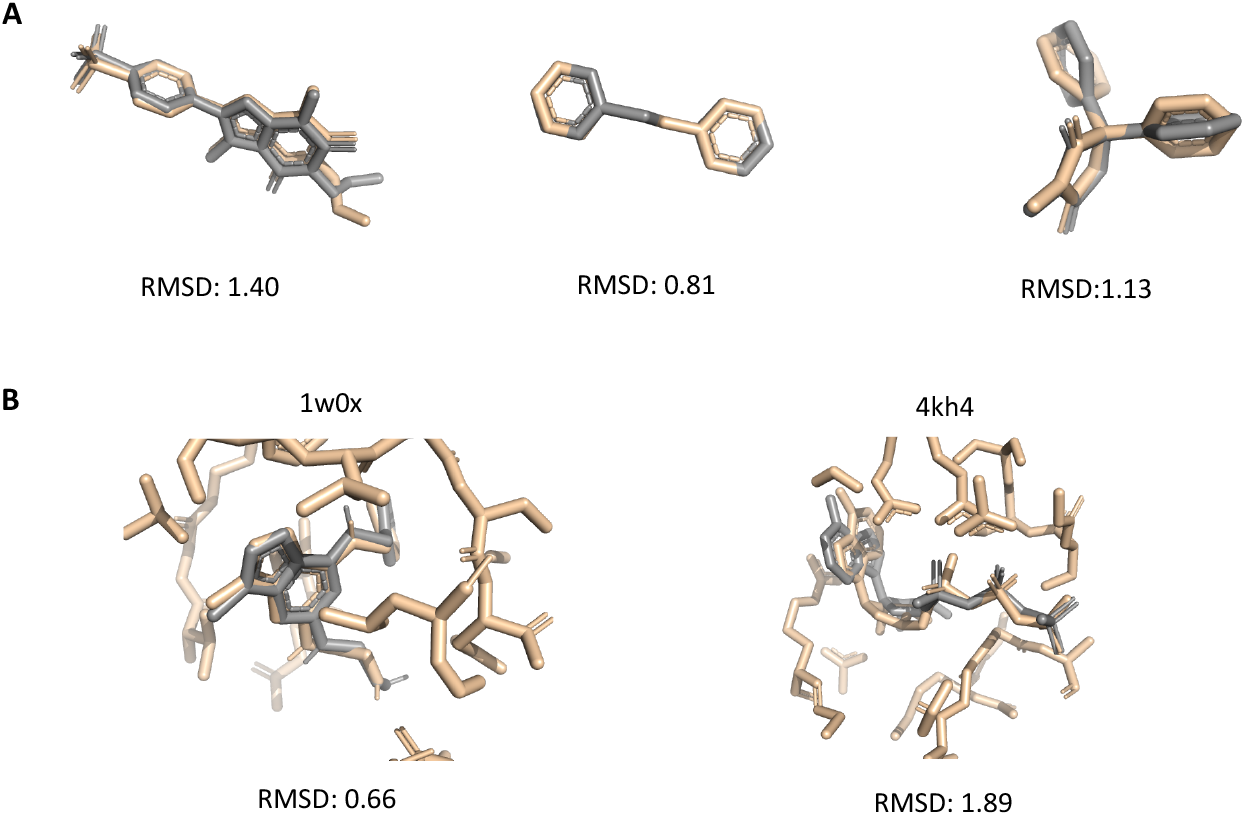
Randomly selected samples from pretrained models on (A) Pubchem and (B) Plinder datasets.

**Figure S2:**
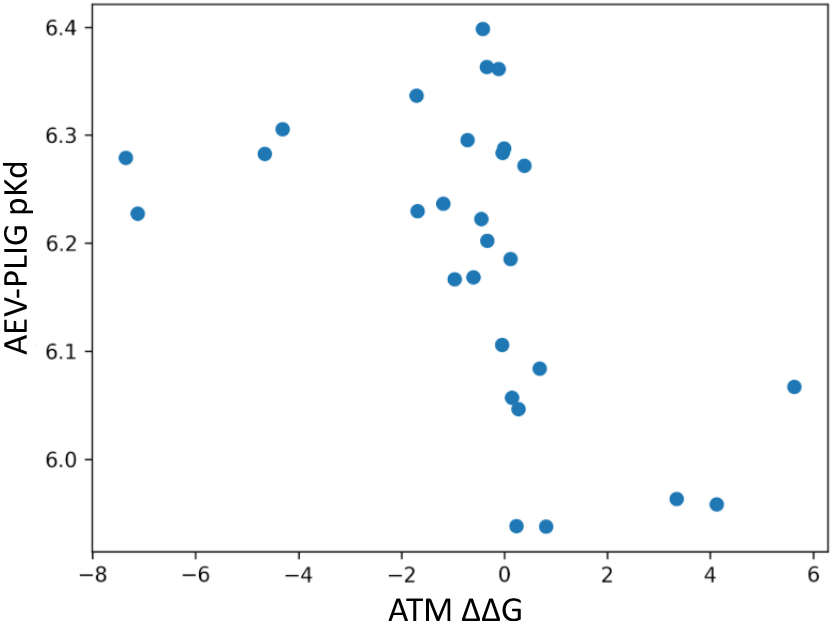
ATM ΔΔ*G* and AEV-PLIG *pK*_*d*_ on PDB 1SSC with a smaller set of ncAAs from CycPeptMPDB [38], exhibiting a Spearman correlation of -0.62 with p < 0.01.

**Figure S3:**
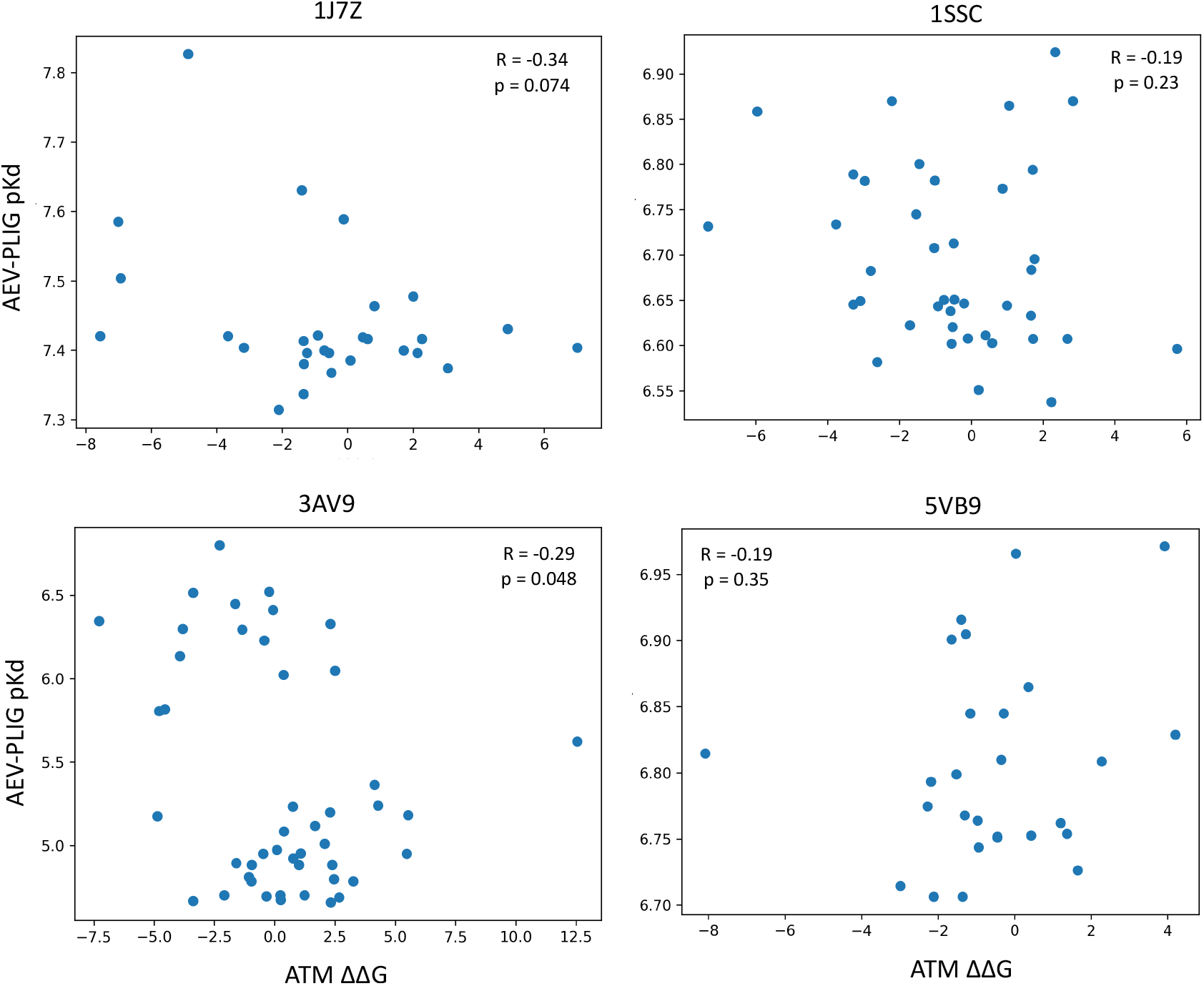
ATM ΔΔ*G* and AEV-PLIG *pK*_*d*_ correlation on four test cases.

**Figure S4:**
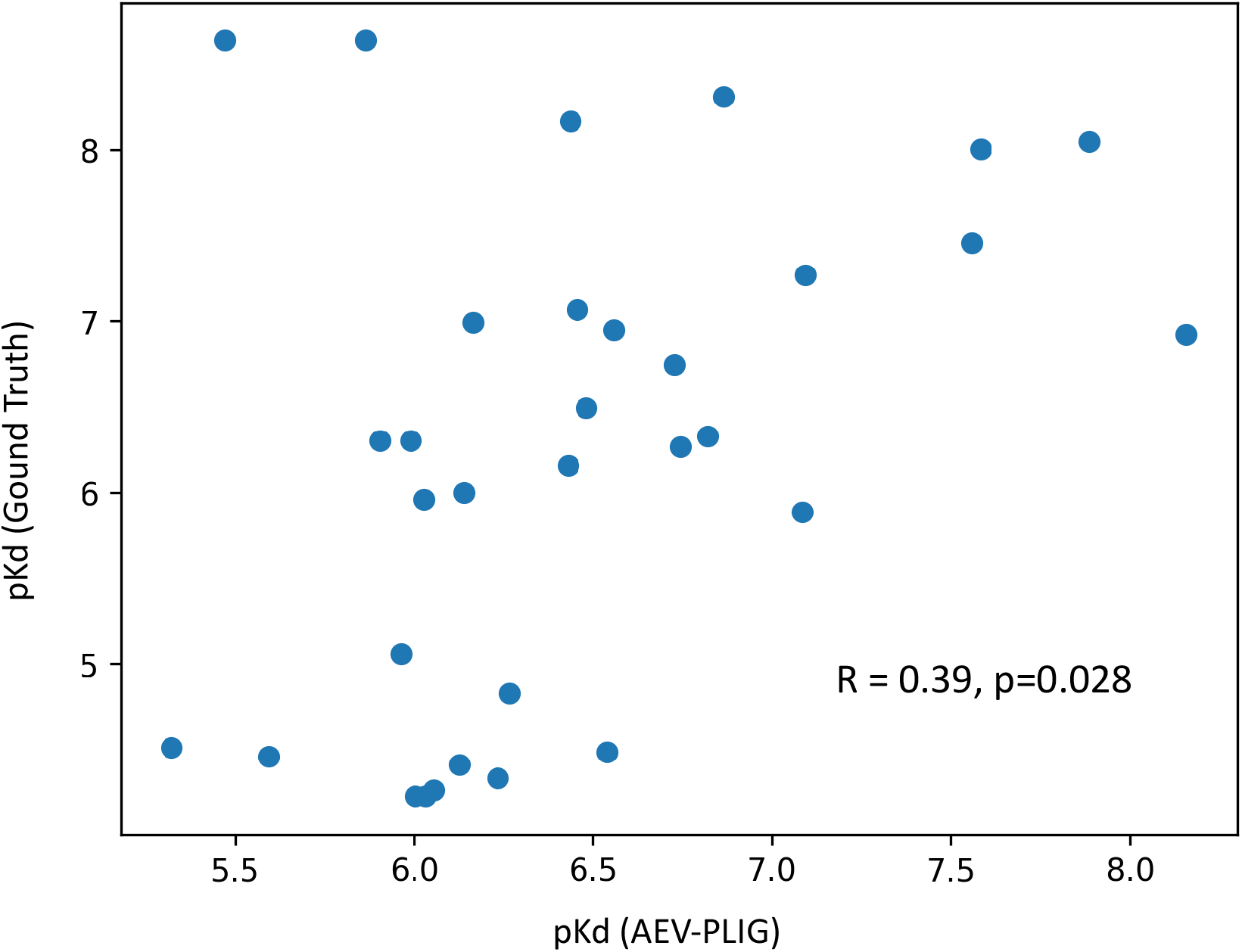
AEV-PLIG predictions vs experimental Kd values on long peptides (20-30aa) in PDB-Bind2020. These peptides are found in the PDBBind-PP dataset and are not in the training set of AEV-PLIG.

**Figure S5:**
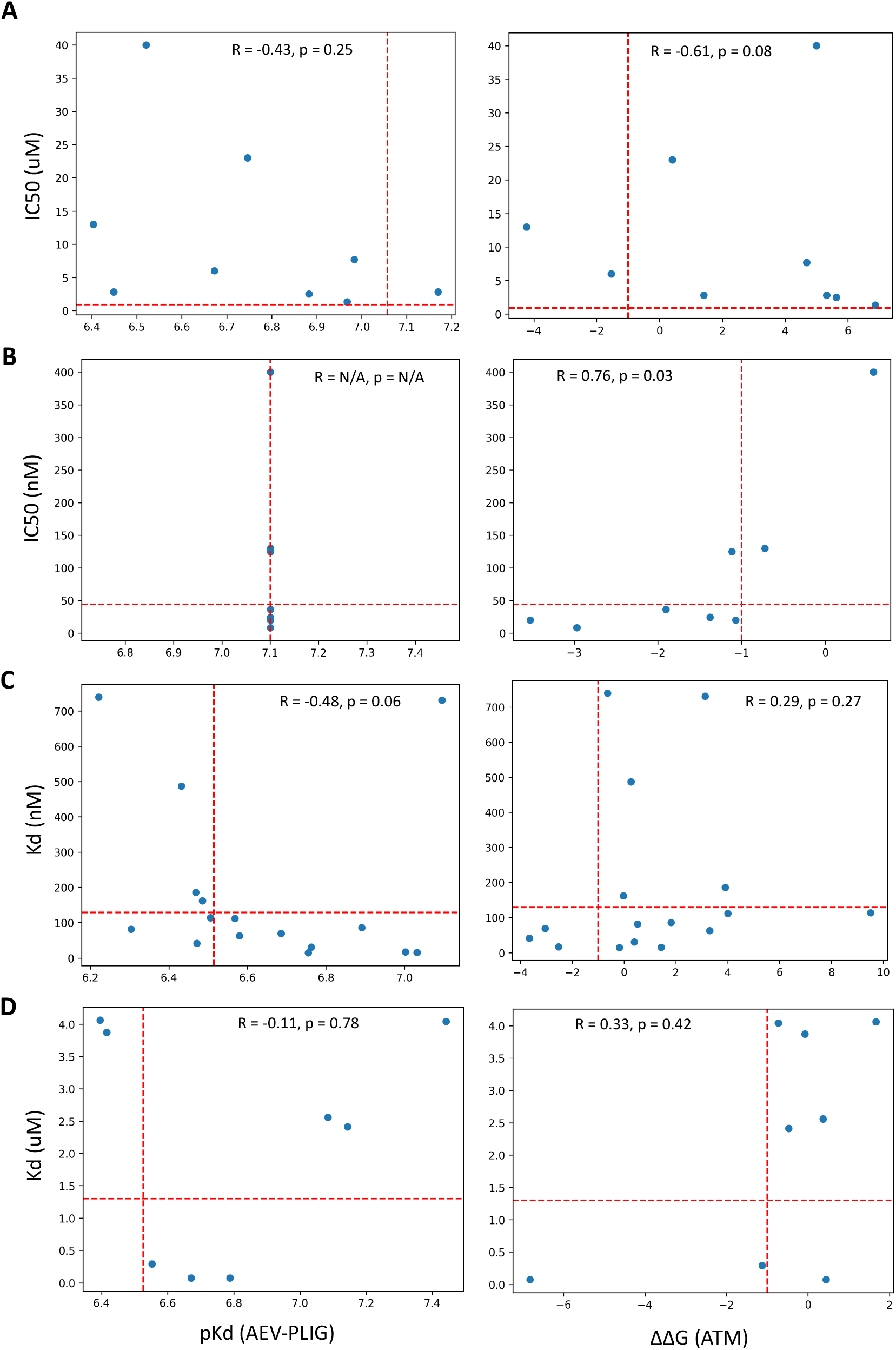
AEV-PLIG (left) and ATM (right) predictions of binding affinity plotted against experimentally measured binding affinities across the four experimental datasets in Table S5, corresponding to (A) Htra1-PDZ, (B) Mdm2-p53, (C) cetuximab-meditope, and (D) Undisclosed. Note that AEV-PLIG appears to predict the same affinity across all variants of Mdm2, perhaps due to

**Figure S6:**
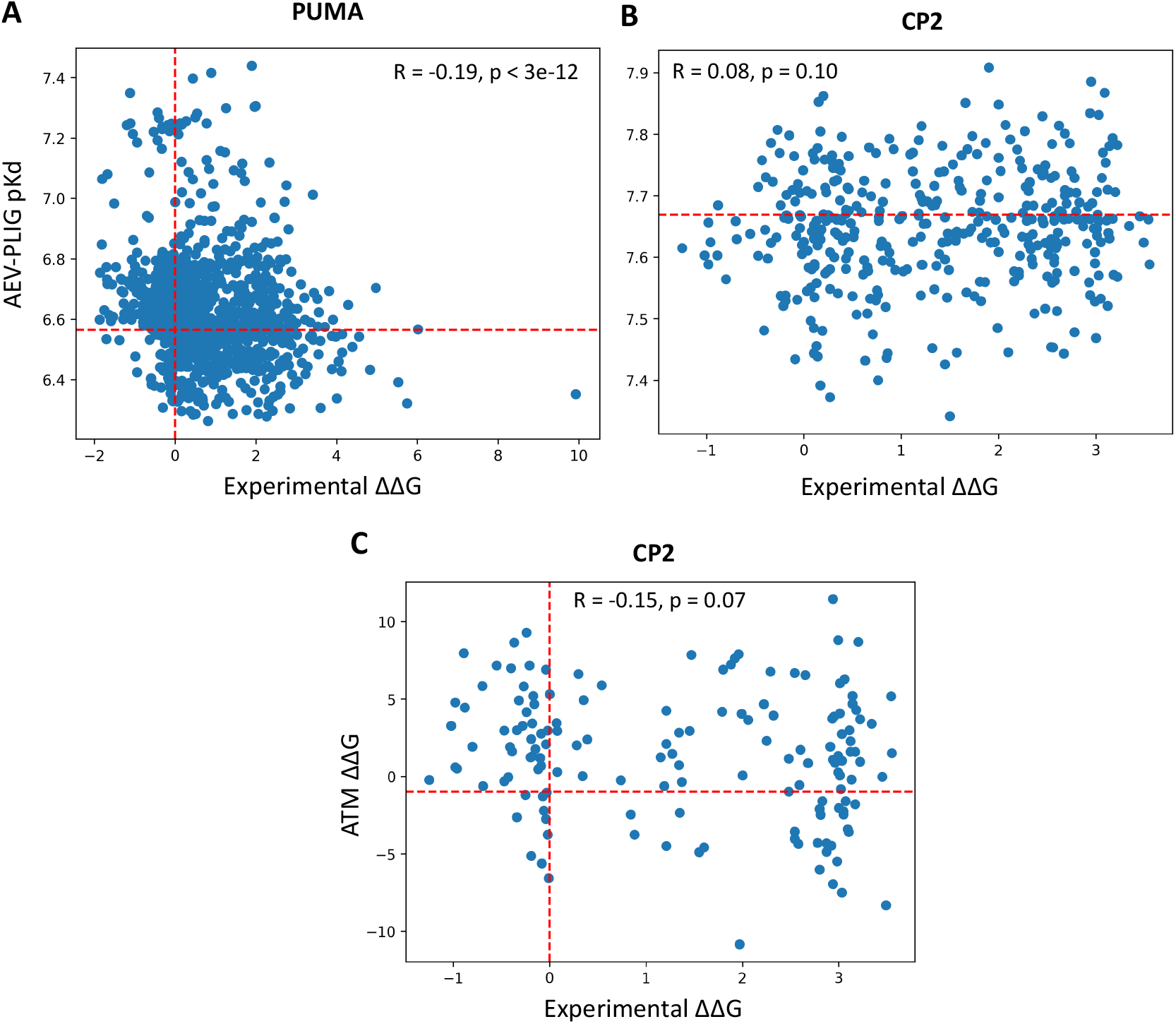
AEV-PLIG and ATM predictions of deep mutational scanning datasets [42] on the peptides PUMA (A) and CP2 (B,C). ATM on the PUMA peptide was unsuccessful due to its length (35aa).

**Table S1:** Number of ncAAs per parent amino acid.

| Parent AA | ncAA Count |
| --- | --- |
| LYS | 111 |
| PHE | 379 |
| PRO | 35 |
| ASN | 9 |
| MET | 7 |
| TRP | 67 |
| ARG | 36 |
| SER | 23 |
| TYR | 63 |
| CYS | 49 |
| GLY | 57 |
| GLU | 17 |
| THR | 17 |
| GLN | 7 |
| ALA | 19 |
| ASP | 10 |
| HIS | 25 |
| LEU | 5 |
| VAL | 3 |
| ILE | 0 |
| **Total** | **979** |

**Table S2:**
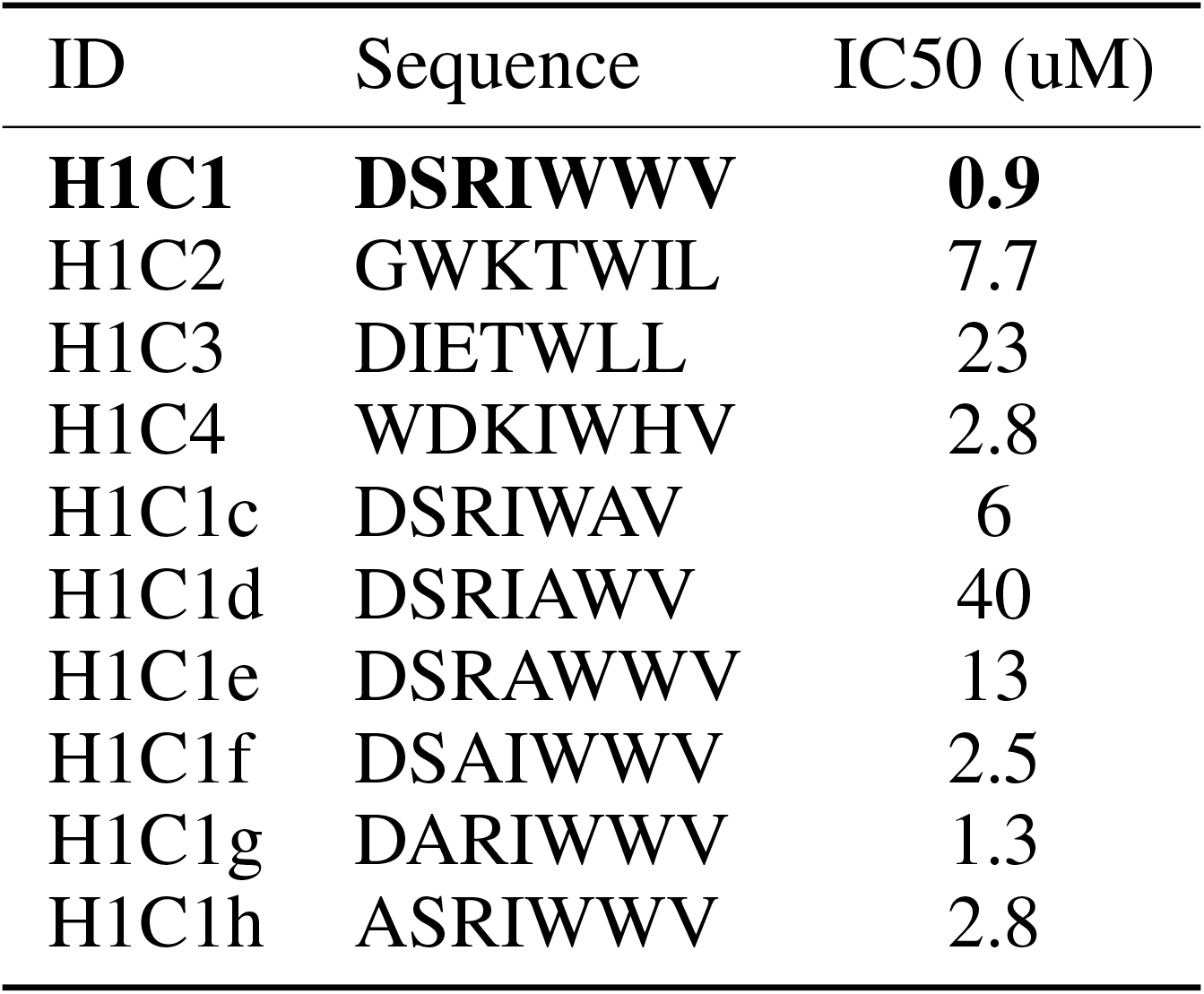
List of variants and corresponding binding affinities for HtrA1 PDZ domain [39]. Bold indicates the reference peptide used for relative binding affinity calculations.

**Table S3:**
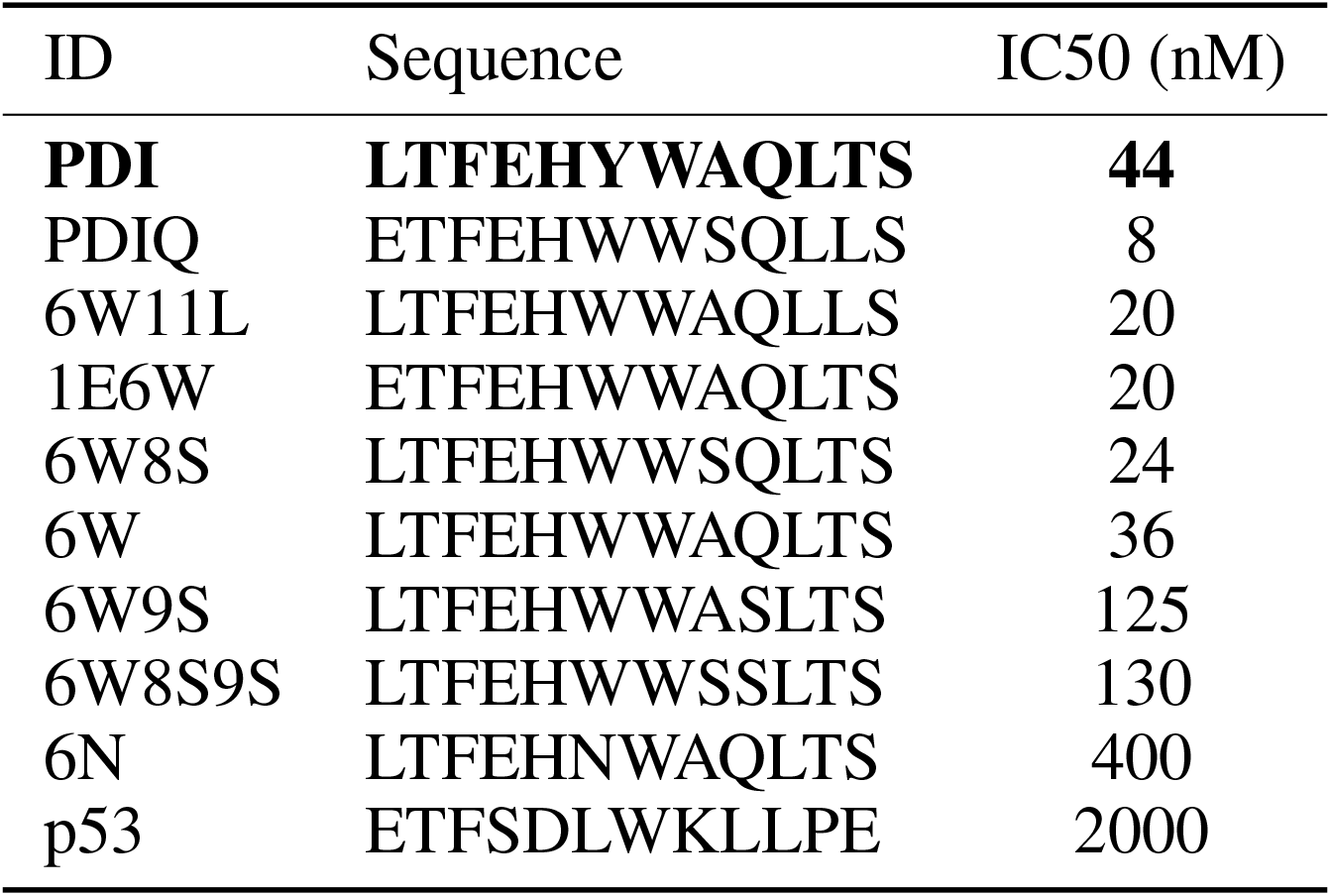
List of variants and corresponding binding affinities for Mdm2-p53 [40]. Bold indicates the reference peptide used for relative binding affinity calculations.

| ID | Sequence | IC50 (nM) |
| --- | --- | --- |
| **PDI** | **LTFEHYWAQLTS** | **44** |
| PDIQ | ETFEHWWSQLLS | 8 |
| 6W11L | LTFEHWWAQLLS | 20 |
| 1E6W | ETFEHWWAQLTS | 20 |
| 6W8S | LTFEHWWSQLTS | 24 |
| 6W | LTFEHWWAQLTS | 36 |
| 6W9S | LTFEHWWASLTS | 125 |
| 6W8S9S | LTFEHWWSSLTS | 130 |
| 6N | LTFEHNWAQLTS | 400 |
| p53 | ETFSDLWKLLPE | 2000 |

**Table S4:** List of variants and corresponding binding affinities for cetuximab-meditope [41]. Bold indicates the reference peptide used for relative binding affinity calculations.

| ID | Sequence | Kd (nM) |
| --- | --- | --- |
| **Md1** | **CQFDLSTRRLKC** | **129.4** |
| Md3 | CQYNLSSRALKC | 739 |
| Md2 | CVWQRWQKSYVC | 731 |
| Q1V | CVFDLSTRRLKC | 111.5 |
| S5G | CQFDLGTRRLKC | 113.9 |
| K10R | CQFDLSTRRLRC | 81.3 |
| Q1V_S5G | CVFDLGTRRLKC | 69.4 |
| Q1V_K10R | CVFDLSTRRLRC | 63.2 |
| S5G_K10R | CQFDLGTRRLRC | 41.1 |
| Q1V_S5G_K10R | CVFDLGTRRLRC | 30.3 |
| Q1V_D3N_S5G_K10R | CVFNLGTRRLRC | 14.5 |
| Q1V_S5G_T6I_K10R | CVFDLGIRRLRC | 16.8 |
| Q1V_D3N_S5G_T6I_K10R | CVFNLGIRRLRC | 15.8 |
| Q1V_S5Y_K10R | CVFDLYTRRLRC | 86.3 |
| Q1V_S5G_T6M_K10R | CVFDLGMRRLRC | 162 |
| Q1V_S5Y_T6M_K10R | CVFDLYMRRLRC | 487 |
| Q1V_D3R_S5G_K10R | CVFRLGTRRLRC | 186 |

**Table S5:** Performance evaluation of AEV-PLIG and ATM binding affinity predictions on four protein-peptide variant datasets.

| Target | Method | Accuracy | Precision | Recall |
| --- | --- | --- | --- | --- |
| HtrA1 [39] | AEV-PLIG | 0.89 | N/A | N/A |
|  | ATM | 0.78 | N/A | N/A |
|  | Both | **1.00** | N/A | N/A |
| Mdm2 [40] | AEV-PLIG | N/A | N/A | N/A |
|  | ATM | **0.67** | **1.00** | **0.67** |
|  | Both | **0.67** | **1.00** | **0.67** |
| Cetuximab [41] | AEV-PLIG | 0.75 | 0.89 | 0.73 |
|  | ATM | **0.80** | 0.50 | **1.00** |
|  | Both | 0.44 | **1.00** | 0.18 |
| Undisclosed<sup>1</sup> | AEV-PLIG | 0.63 | 0.50 | **1.00** |
|  | ATM | **0.88** | **1.00** | 0.67 |
|  | Both | **0.88** | **1.00** | 0.67 |

<sup>1</sup> Due to conflict of interest.

## Notes

### Competing Interest Statement

J.S.L. is a consultant for a stealth startup, and PMK is a co-founder and consultant to several biotechnology companies, including Fable Therapeutics and TBG Therapeutics.

### Funding

- Canadian Institutes of Health Research (CIHR) Project Grant: PJT-166008

